# Sparse Representation for High-dimensional Multiclass Microarray Data Classification

**DOI:** 10.1101/2023.12.19.572302

**Authors:** Maliheh Miri, Mohammad Taghi Sadeghi, Vahid Abootalebi

**Affiliations:** Department of Electrical Engineering, Yazd University, Yazd, Iran

**Author notes:** Corresponding author. E-mail address (Maliheh Miri).

**Keywords:** Sparse Representation, Sparse Representation-based Classifiers, Microarray Data, Gene Expression, Tumor Classification

## Abstract

Sparse representation of signals has achieved satisfactory results in classification applications compared to the conventional methods. Microarray data, which are obtained from monitoring the expression levels of thousands of genes simultaneously, have very high dimensions in relation to the small number of samples. This has led to the weaknesses of state-of-the-art classifiers to cope with the microarray data classification problem. The ability of the sparse representation to represent the signals as a linear combination of a small number of training data and to provide a brief description of signals led to reducing computational complexity as well as increasing classification accuracy in many applications. Using all training samples in the dictionary imposes a high computational burden on the sparse coding stage of high dimensional data. Proposed solutions to solve this problem can be roughly divided into two categories: selection of a subset of training data using different criteria, or learning a concise dictionary. Another important factor in increasing the speed and accuracy of a sparse representation-based classifier is the algorithm which is used to solve the related ℓ^1^ –norm minimization problem. In this paper, different sparse representation-based classification methods are investigated in order to tackle the problem of 14-Tumors microarray data classification. Our experimental results show that good performances are obtained by selecting a subset of the original atoms and learning the associated dictionary. Also, using SL0 sparse coding algorithm increases speed, and in most cases, accuracy of the classifiers.

## 1. Introduction

Sparse Representation (SR) of signals has great importance in different signal processing applications and many studies have been conducted in this area [1, 2]. In many pattern recognition and computer vision applications, we are dealing with high-dimensional data, such as face images or microarray data [3].

In 2009, Wright proposed the use of sparse representation of signals in classification problems. This method, called sparse representation-based classification (SRC), has led to satisfactory results, especially in face recognition systems [4]. The main idea behind SRC is that each face image can be represented as a sparse linear combination of the other faces, such that the largest coefficients in this linear combination belong to the images that are in the same class with the input image. Therefore, by using these larger coefficients and by resetting the other coefficients to zero, a reasonable reconstruction of the input image can be achieved. Euclidean distance metric is used for determining the similarity between the input image and reconstructed images by different classes. Finally, the test data belonged to the class resulting in a minimal reconstruction error.

Advances in molecular biology and microarray technology have made it possible to monitor the expression levels of thousands of genes (gene activity) simultaneously, leading to the production of massive amounts of microarray data [5]. These data play an important role in the diagnosis and classification of various cancer tissues, but their most challenging aspects can be identified as their very high dimensions compared to the small sample size, which makes design of appropriate classifiers difficult. Many attempts have been made in order to find high accuracy and low computational complexity classifiers. However, most of these studies focused on the data with a small number of classes (two or three different tumor types), and applied methods such as Linear Discriminant Analysis (LDA), Neural Networks, Clustering, Nearest Neighbour (NN), and Support Vector Machine (SVM), Decision tree, and Random Forest [5-12]. In this study, the 14-Tumors database is taken under examination. This database includes information on fourteen different tumor types that have often been studied from the perspective of feature selection. Among the classification works that have been conducted on this database, the method presented in [5] can be noted where the classification problem was converted to a binary classification problem, and different classifiers were used for this purpose of which the best performance was obtained from the SVM classifier. In the binary classifiers, classification procedure is performed using One vs. All (OVA) and One vs. One (OVO) methods that respectively use *c* (*c* −1) / 2 and *c* hyperplanes to separate the classes (*c* is the number of classes). In [6] and [7], the 14 tumors data were classified by using neural network based methods. For example, in [5], two consecutive Artificial Neural Networks (ANN) classifiers are used for classification. The first classifier selects two classes that the test data most likely belongs to, and the second one makes the final decision among the selected classes. A fuzzy classifier was presented in [8] that uses fuzzy rules for classification. In [9], a sparse representation based method has been applied in order to classify tumor data. It has been shown that their proposed method outperforms various extensions of the SVM classifier.

Despite the wide research in this area [13-16], the classification performance of the reported classifiers is not meaningful in clinical diagnosis. Inspired by impressive results of using sparse representation in processing of gene expression data [17-21], in this paper, several sparse representation-based classification approaches are considered to overcome the problem of 14-Tumors classification. In order to obtain a more accurate evaluation, three different algorithms for the sparse coding stage are utilized, each of which calculates the sparse representation using a different approach.

This paper is structured as follows: Section 2 presents an introduction to the sparse representation. Some explanations on conventional sparse coding algorithms is then given in Section 3. In Section 4, the adopted sparse representation-based classifiers are reviewed. In Section 5, a brief description of the used database is given and the experimental results are reported. Finally, discussions and conclusions are presented in sections 6 and 7.

## 2. Sparse Representation Theorem

In recent years, sparse representation has become a powerful tool for signal representation and compression. According to this representation, a signal can be approximated as a sparse linear combination of a set of training data [2]. Suppose **Y** = [**Y**_1_, …, **Y**_*c*_ ] is composed by concatenating all the training data so that **Y** _1_, …, **Y** _*c*_ contains the samples of classes 1 to c respectively. Therefore, a linear combination of the training data can be found by a system of linear equations as **y = Ys**, in which **Y** is the dictionary matrix and each of its column is called an atom.

If the number of equations is less than that of number of variables (the number of **Y** columns is greater than that of its rows), the system is under-determined and has countless solutions. Imposing sparsity constraint on the solutions leads to a unique solution. Since *ℓ*^0^ -norm of a vector represents the number of its non-zero elements, minimizing the *ℓ*^0^ -norm leads to the imposition of sparsity constraint. Therefore, the following equation is used in order to determine the sparse representation of an input signal on the associated training set:

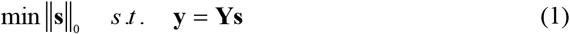

where, **y** is input signal, **s** is the coefficient vector of the linear combination, ‖ **s** ‖_0_ and shows the number of its non-zero elements. **s** is k-sparse if it has at most k nonzero elements.

Assuming *N* _*i*_ training data belonging to the i-th class is to be placed in the columns of the **Y** matrix as 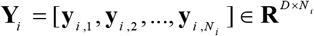, if **y** ∈ **R**^*D*^ is the input data belonging to the i-th class, according to the sparse representation it can be represented by merely using the **Y**_*i*_ columns as follows:

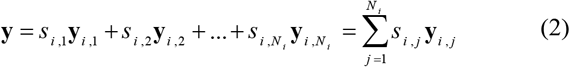

in which, *s*_*i, j*_ is scalar. This is the idea behind sparse representation-based classification (SRC) [4].

Since *ℓ*^0^ -norm is non-convex, determining the sparse representation using equation (1) is NP-hard, and its solution depends on the combinatorial search that is impossible in high dimensions [2]. To solve this problem, several methods have been proposed, some of which will be discussed in the following section.

## 3. Sparse Coding Algorithms

A conventional method for solving linear systems by assuming the sparsity of solution is approximation of *ℓ*^0^ - norm with a higher-order norm that is easier to minimize. Therefore, in order to determine the sparse representation of signal, the following equation can be used in which the sparsity constraint is provided by *ℓ*^1^ -norm minimization:

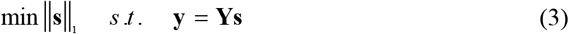

Since *ℓ*^1^ -norm is convex, its minimization can be calculated using convex optimization methods; it also can be expressed as a linear problem and be solved using linear programming methods such as Basis Pursuit (BP) [22]. It has been proved that if the number of non-zero elements in sparse solution is less than a certain number depending on the correlation between dictionary atoms, the answer to equation (3) is equal to the *ℓ*^0^ -norm minimization solution [2]. One of the main disadvantages of the BP method is its high computational complexity. As its implementation is time consuming, it is often replaced by other methods.

One of the fastest algorithms to determine the sparse representation is Matching Pursuit (MP) [23]. This method acts greedily and repetitively so that only a coefficient is determined at each step. If **s** be a k-sparse vector, then **y** can be written as a linear combination of k atoms of the dictionary. In this method, firstly the columns of dictionary which are present in the linear combination are detected, and then related coefficients to these columns, the non-zero elements of **s**, are calculated by solving a least square equation. Therefore, at each step, one atom of the dictionary that is most similar to the test sample is considered an active member of the linear combination and its related coefficient is calculated. Subtracting the product of this 1-sparse approximation and dictionary from the test sample is considered a residual, and the above steps are repeated. The sum of the 1-sparse approximations obtained at each step with previous approximations is considered a new approximation, and this process continues until either a certain number of steps are completed or the error is less than a certain amount. Since this method requires a simple search at each step, it is usually very fast, but because of its greedy nature there is no guarantee that the final solution will be the same as the sparse solution.

At any stage in Orthogonal Matching Pursuit (OMP), which is an extension of the MP method, related coefficients to active columns of the dictionary are selected independently from the previous results, and the previous results are merely used for locating non-zero elements [24].

One of the other methods developed in order to find the sparse representation is Smoothed *ℓ*^0^ -norm (SL0) [25]. In this method, instead of replacing the *ℓ*^0^ –norm with a higher-order norm, *ℓ*^0^ -norm itself is minimized. In Figure 1 (a), one variable *ℓ*^0^ -norm function is shown, which is zero in zero place and 1 in the other places. As can be seen in this figure, *ℓ*^0^ -norm function is not continuous. Therefore, SL0 uses a smooth function that provides an approximation of the *ℓ*^0^ -norm and can be optimized easily. In Figure 1 (b), a smooth function has been depicted acting somewhat similar to the *ℓ*^0^ –norm (symmetric Gaussian function). If this function is named as *f* _1_ (*s*), the new optimization problem is as follows:

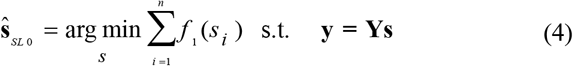

If *f* _1_ (*s*) is not smooth enough, a lot of local minimum possibly will exist. For this reason, a very smooth function (Gaussian with large variance) is selected at first, which has only a local minimum and after finding the optimum location, this value is used to initialize the function that is closer to the *ℓ*^0^ -norm (reduced variance). Therefore, the previous result is used at each step and the function will be closer to the *ℓ*^0^ -norm. It is proved that with proper selection of variances, obtained solution in infinite (zero variance) converges to the *ℓ*^0^ –norm solution.

**Fig. 1.**
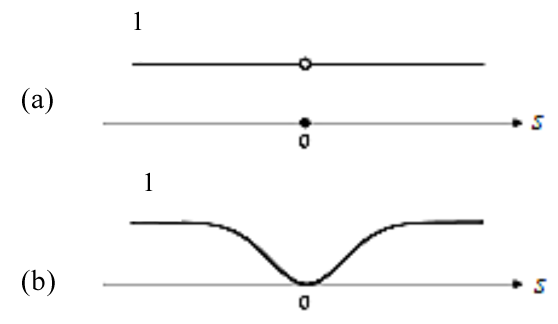
*ℓ*^0^ -norm Function (a), Smooth Function (b).

One of the advantages of the SL0 method is its high speed. It is stated that its convergence speed is two to three times faster than the usual algorithms for solving linear problems. In addition, since this method attempts to minimize the *ℓ*^0^ -norm, it is expected that more complicated sparse problems will be solvable using this algorithm [25]. Some of the other algorithms that can be used to calculate the sparse representation are Homotopy, GPSR^1^, FISTA^2^, CoSaMP^3^, and IHT^4^ [26-30].

## 4. Sparse Representation-based Classifiers

In this section, the most important sparse representation based classifiers are reviewed.

### 4.1. Basic Sparse Representation based Classification (SRC)

This method involves two stages: First, the test sample is coded sparsely using a dictionary matrix which is formed by combining all training data from all classes:

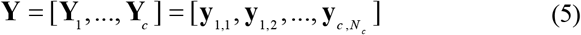

in which **Y** is a sub-matrix that contains the training samples of the i-th class. The representation is obtained using equation (3) and the discussed algorithms.

SRC classifies the test sample by evaluating the similarity between the sample and the training data in each class.

For this purpose, *δ* _*i*_: **R**^*N*^ →**R**^*N*^ function is defined as:

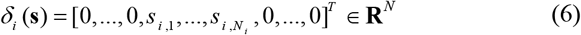

By applying this function to the **s** vector, *N*_*i*_ elements of **s** corresponding to the i-th class samples are preserved and the others become zero.

Therefore, ŷ _*i*_ = **y** *δ* _*i*_ (**s**) expresses **y** as a linear combination of the samples in i-th class. By applying this function on the representation **s** and reconstructing **y** using samples belonging to each class, the test sample will be assigned to the class that minimizes the following residual [4]:

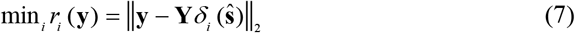

### 4.2. Collaborative Representation based Classification (CRC)

It has been generally accepted that sparsity plays a key role in the success of the SRC method and the role of effective participation of training samples from different classes in representing the input sample is almost ignored. In [31], it has shown that if the number of training samples within the dictionary is large in comparison with the data dimension, *ℓ*^2^ -norm has an acceptable performance compared to the *ℓ*^1^ -norm. Therefore, the following equation with low computational complexity has been used to obtain the representation of the input sample by the participation of all training samples:

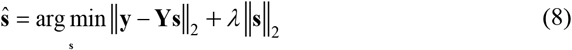

where *λ* is a scalar value. In fact, the *ℓ*^1^ –norm minimization constraint has been replaced by *ℓ*^2^ -norm. *ℓ*^2^ -norm minimization not only imposes a certain amount of sparsity on the coefficients, but also makes the least square solution stable. The solution of the above equation using the least square will be as follows:

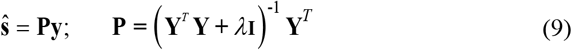

where, **I** is the identity matrix. Since **P** is independent of the input data, it can be pre-calculated and by entering the test sample, it is mapped onto **P** which leads to much lower complexity and higher speed of the CRC compared to the SRC. The general equation of collaborative representation classification is as follows:

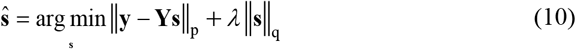

The more restrictive the constraints on the reconstruction residual, the more guarantees the stability of the classifier. Furthermore, *ℓ*^1^ -norm or *ℓ*^2^ –norm minimization constraints on the sparse coding coefficients is proportional to discriminatory ability of the representation [31]. Therefore, the SRC method can be considered a special case of CRC, by putting p = 2 and q = 1.

### 4.3. Metasample based Sparse Representation Classification (MSRC)

The main idea in metasample based SRC is the representation of test sample as a linear combination of metasamples which are extracted from the training samples [21]. A metasample is a linear combination of samples that contains their intrinsic properties. Mathematically, the matrix of training samples (original dictionary) can be decomposed into two matrices as follows:

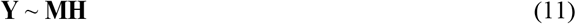

in which, **M** is a *D* × *P* matrix in which each column defines a metasample, and **H** is a *P* × *N* matrix where each column represents the relationship between metasamples and their corresponding samples in the main dictionary.

Several methods are proposed for extracting metasamples from the original samples. In [32], metasamples are extracted using Non-negative Matrix Factorization (NMF) method. In [33], Singular Value Decomposition (SVD) is used for this purpose, where the eigenvalues of the training samples matrix are calculated and sorted from the largest to the smallest. Then, instead of the original samples, a number of their corresponding eigenvectors are used which provide a general picture of the dynamics of the data.

In order to use metasamples in classification via sparse representation, the metasamples are determined for each class, separately, as follows:

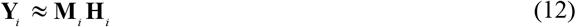

where **M**_*i*_ is a *D* × *P*_*i*_ matrix each column of which represents a metasample of the i-th class. The number of selected metasamples for each class can be determined experimentally or by cross validation. Matrix **M** is composed by concatenating the calculated metasamples for each class, as follows:

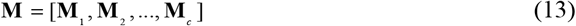

By entering a new test sample, its sparse representation on the metasamples matrix is calculated.

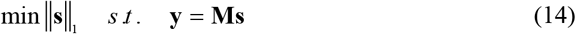

Finally, test label can be determined using the method presented in the SRC. Therefore, the main difference between the SRC and MSRC is in dictionary atoms which unlike SRC that uses original training samples in the dictionary, MSRC puts extracted metasamples for each class in the dictionary. In [19], a Parameter Free MSRC method (PFMSRC) was proposed which addresses optimal selection of the number of metasamples and the sparse penalty factor by a weighting strategy.

Li et al. presented a tumor classification method called MRSRC (Maxdenominator Reweighted Sparse Representation Classification) [18] in which, first, a set of meta-genes is extracted from the training samples and then a reweighted *ℓ*^1^ regularization method is used to obtain the sparse representation coefficients. Finally, a maxdenominator residual error function is utilized for classification. In their study, binary/multiclass microarray data sets were used to compare the performance of different classification methods. The obtained results show that sparse representation-based methods (SRC, MSRC and MRSRC) outperform the other methods in most experiments, which indicates the capability of these methods in small sample size classification problems.

### 4.4. K Nearest Neighbour based SRC (KNN-SRC)

In data with high dimensions or large number of samples, using the entire training samples in the dictionary imposes high computational burden on the sparse coding stage and decreases the speed of the SRC algorithm.

Some methods speed up the classification process by choosing a subset of training data to solve *ℓ*^1^ –norm minimization problem. In these methods, a subset of atoms from the original dictionary which meets a predetermined criterion, according to the application, is selected to form the final dictionary. For example, in [34] it is shown that imposing sparsity constraint on the atoms of dictionary leads to sparser representations and consequently better results in speech denoising. Also, in [35], the classification accuracy has improved by selecting sparse atoms in order to form a concise dictionary.

In KNN-SRC method, computational complexity is significantly reduced by selecting *K* nearest neighbours to the test sample and by including the selected samples in the classification process [36].

In this method, the *K* nearest neighbours to the test sample are found by applying Euclidean distance. The test sample is then sparsely represented by these *K* neighbours. If 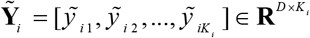 contains *K* _*i*_ neighbours from i-th class, so that 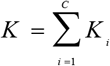, and matrix 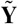 includes all *K* nearest neighbours, the sparse representation of **y** is calculated as follows:

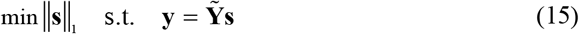

Since *K* ≪ *N*, **y** never would be quite equal to 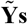. Therefore, the following equation is used for calculating the sparse representation:

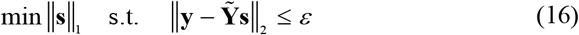

After determining the representation, the classification process is performed similar to the SRC method.

### 4.5. Linearly Approximated SRC (LASRC)

It is shown in [37] that effective training samples in the sparse representation of signal are not necessarily the nearest samples which are determined by Euclidean distance metric. Therefore, attempt is made to predetermine the training samples which will be chosen by *ℓ*^1^ -norm minimization in order to form a concise dictionary.

The objective function of the Lagrangian formulation of equation (3) can be expanded as follows:

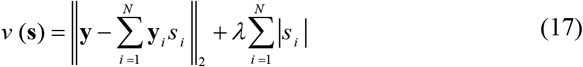

Suppose **s** is k-sparse. Therefore, for any *I* Where *s* _*i*_ = 0, then = ‖ **y** _*i*_ s _*i*_ ‖ = 0, | *s* _*i*_ | = 0. Thus, the above equation is rewritten as follows:

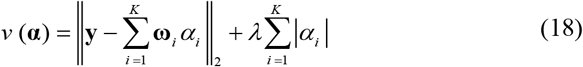

where **ω** represents a column of **Ω**. The new dictionary **Ω** only contains atoms of the original dictionary which are corresponding to the non-zero elements of **s**. Therefore, the *ℓ*^1^ -norm minimization using **Ω** and ***α*** coefficients vector will be as follows:

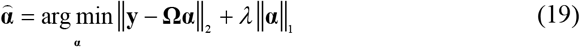

Since finding precise location of effective atoms in the sparse representation is a difficult task, these atoms are approximated using *ℓ*^2^ -norm minimization. Although the solution of *ℓ*^2^ -norm minimization is dense and has a relatively high number of non-zero elements, in [37] it is shown that the largest peaks occur in the similar locations to the *ℓ*^1^ -norm minimization solution. Therefore, the solution of the *ℓ*^2^ -norm minimization can be considered an initialization point for calculating the sparse representation using the *ℓ*^1^ -norm minimization.

As discussed in the CRC method (section 4.2.), replacing *ℓ*^1^ -norm by *ℓ*^2^ -norm in sparse coding equation leads to least square equation which has much higher speed compared to the *ℓ*^1^ -norm minimization. Therefore, in this method, by thresholding the *ℓ*^2^ –norm minimization solution, the training samples which have the largest magnitudes are selected for initializing the *ℓ*^1^ - norm minimization. This method is called Linearly Approximated SRC (LASRC) which combines the speed of the CRC method and the robustness of the SRC method and has achieved significant results in face verification systems [37,38].

### 4.6. Sparse Hierarchical Classification (SSC-SRC)

In [39], a hierarchical classification method using sparse representation has been presented. In this method, a subset of the training samples which are in the same cluster with the test sample is used to form the final dictionary. This method which uses concise dictionary has achieved a higher classification accuracy than the SRC.

In clustering stage, sparse subspace clustering (SSC) method is used which separates different clusters of training samples using their sparse representations, and classification stage has been carried out using the SRC. In the next section, following a brief review of the SSC method, this classification approach is explained.

#### 4.6.1. Sparse Subspace Clustering (SSC)

Suppose 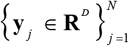 are data from *n* independent linear subspaces 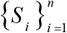 with 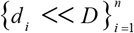 dimensions, and 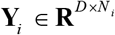 are *N*_*i*_ data belonging to i-th subspace in which 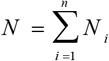 is the total number of data. Matrix **Y** is formed as follows:

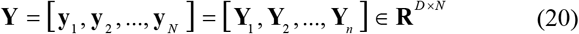

If **y**_*i*_ be a sample from *S* _*i*_ subspace, it can be written as a linear combination of *d* _*i*_ samples in this subspace. In other words, **y** has a *d* -sparse representation that can be determined from equation (3) where for *j* ≠ *i*, **s** ≠ 0 and **s** _*j*_= 0 [40].

For data clustering using obtained sparse representations, spectral clustering is used. In this method by using local information about each sample, similarities are defined between each pair of samples. Then, a clustering procedure is performed using the similarity matrix and dividing data into several groups so that samples in each group are similar to each other and are different from those of other groups.

If **Y**_*î*_ ∈ **R** ^*D* × *N*−1^ is the matrix obtained by elimination i-th column of **Y**, **y**_*i*_ has a sparse representation on **Y**_*î*_ which can be calculated from the following equation:

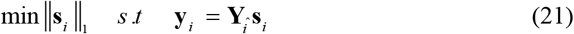

Desired solution **s**_*i*_ ∈ **R**^*N* −1^ is a vector which its non-zero elements are corresponded to the columns of **Y**_*î*_ that are located in the same subspace with **y**_*i*_. After solving the above equation for *i* = 1, …, *N*, coefficients matrix **S** = [**ŝ** _1_, **ŝ** _2_, …, **ŝ** _*N*_] is formed in which, **ŝ** _*I*_ is obtained by putting 0 in i-th row of **s** _*i*_. By using this matrix, graph *G* = (*V, E*) is defined so that vertices *V* are corresponded to *N* data. An edge (*v*_*i*_, *v*_*j*_) ∈ *E* will be exist if **y** _*j*_ be present in the sparse representation of **y**_*i*_. If **y** _*i*_ can be written as a linear combination of a number of samples in a subspace including **y** _*j*_, then **y** _*j*_ can also be written as a linear combination of the samples in the same subspace including **y** _*i*_, so the adjacency matrix of the graph will be as 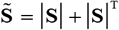.

After construction of the graph, it is expected that the vertices which are related to the same subspace form a connected set in the graph, while the vertices from different subspaces have no common edge. Therefore, 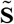 has a block diagonal form as follows:

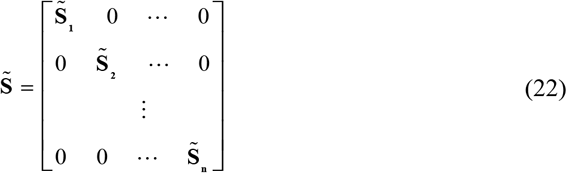

Then, the Laplacian matrix of the similarity graph is formed as 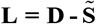 in which **D** ∈ **R**^*N*×*N*^ is a diagonal matrix calculated as 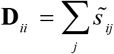. Finally, data clustering is achieved by applying k-means algorithm to n eigenvectors corresponding to n smaller eigenvalues of the Laplacian matrix [40].

#### 4.6.2. SSC-SRC

In this method, by combining sparse representation based clustering and classification methods, classification process is made hierarchically. The main idea behind this method is data clustering and its subsequent classification on each cluster. In other words, after determining the sparse representation of the test sample on the training samples, instead of reconstruction of the signal on the all training samples, the test sample is merely reconstructed on its most similar training samples which are identified in the clustering stage [39].

In SSC-SRC, training samples are clustered in the training phase using SSC algorithm. For this purpose, sparse representation of each training sample on the other samples is achieved using equation (21). After determining the adjacency matrix of the graph and the Laplacian matrix, as described in section 4.6.1., clusters are identified.

In test phase, by entering a test sample into the system, first, its sparse representation on the training samples is achieved. Then, in the reconstruction phase, in each step, sparse coefficients corresponding to one cluster are kept and the remaining coefficients will set to zero. By finding minimum reconstruction error, the test sample’s cluster is detected. In the next stage, in order to determine the test label, sparse representation of the test sample is determined only on the samples belonging to its related cluster which has been identified in the previous stage. Therefore, the final decision will be taken on a smaller subset. In Figure 2 this classification procedure is shown briefly.

**Fig. 2.**
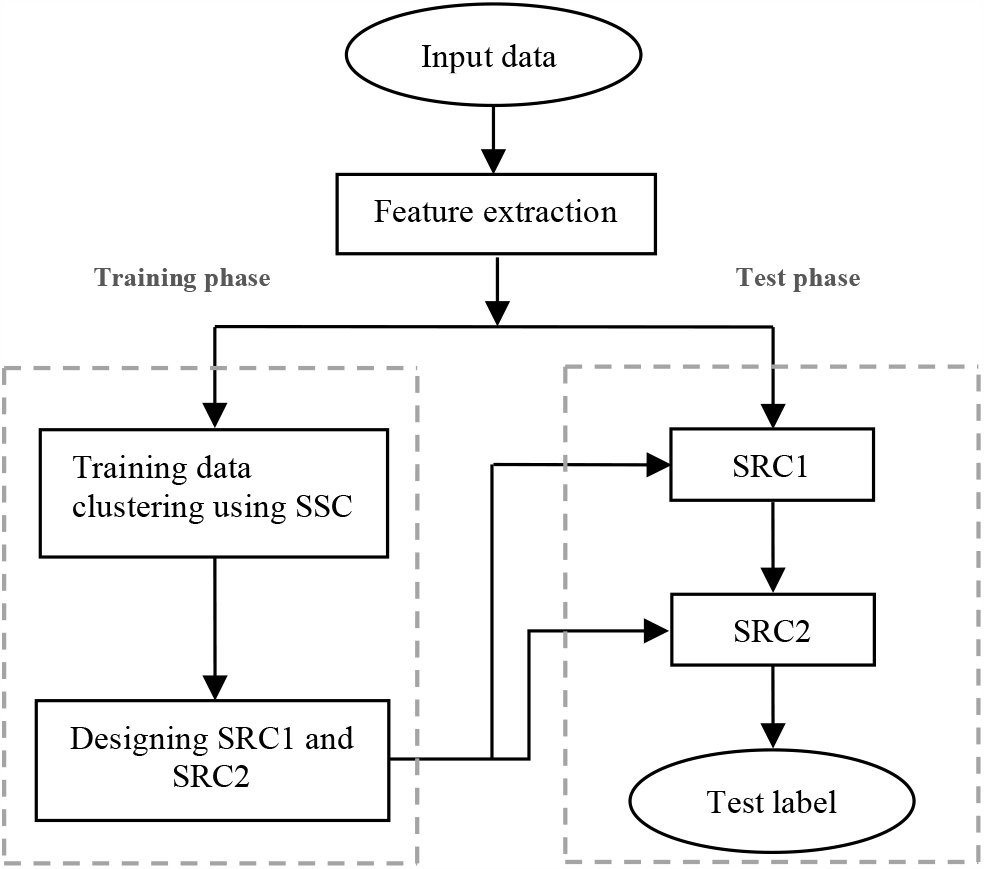
SSC-SRC Block Diagram [39].

## 5. Experiments

In this section, the database used in our experiments is introduced and the simulation results are presented and analysed.

### 5.1. Data

The database we used in our experiments to evaluate the performance of the reviewed classifiers is 14-Tumors database [5]. This database involves 198 samples which contain 16,063 genes (features) of 14 types of tumor, among which 144 samples have been used for training and 54 samples for test. The given value for each gene in each sample is the gene activity, i.e. the gene expression. General characteristics of this dataset are given in Table 1.It can be seen that the number of samples in each class is much lower than their dimensions (16,063).

**TABLE I.** data characteristics.

| Tumor types | Number of training samples | Number of test samples |
| --- | --- | --- |
| Breast | 8 | 4 |
| Prostate | 8 | 6 |
| Lung | 8 | 4 |
| Colorectal | 8 | 4 |
| Lymphoma | 16 | 6 |
| Bladder | 8 | 3 |
| Melanoma | 8 | 2 |
| Uterus | 8 | 2 |
| Leukemia | 24 | 6 |
| Renal | 8 | 3 |
| Pancreas | 8 | 3 |
| Ovarian | 8 | 4 |
| Mesothelioma | 8 | 3 |
| Central Nerve System (CNS) | 16 | 4 |

### 5.2. Simulation Results

The classification of the mentioned database is a challenging problem and most of the commonly used classification methods lead to poor results. In [9], the associated classification results using various extensions of the SVM classifier including the OVR^5^, OVO^6^, DAG^7^, WW^8^ and CS^9^ methods have been reported. In their experiments, polynomial and Radial Basis Function (RBF) kernels were used and each kernel parameter was tuned using the cross validation technique. The lowest obtained error is about 24%. In the following, results of applying different sparse representation-based classifiers are presented.

High dimensionality of data not only increases memory requirements and computational complexity of algorithms but also decreases the efficiency of algorithms because of noise and small number of samples (compared to the dimensions)—inferred as curse of dimensionality. In order to determine the optimal value of reduced dimension using PCA, the 10-fold cross validation technique was applied. For this purpose, training samples were divided into ten folds. Then at each step, one fold was regarded as test data and the remaining nine folds were considered training data. For different values of data dimension, the classification procedure was repeated. The results of these experiments within the framework of the SRC method along with the OMP algorithm are shown in Figure 3.

**Fig. 3.**
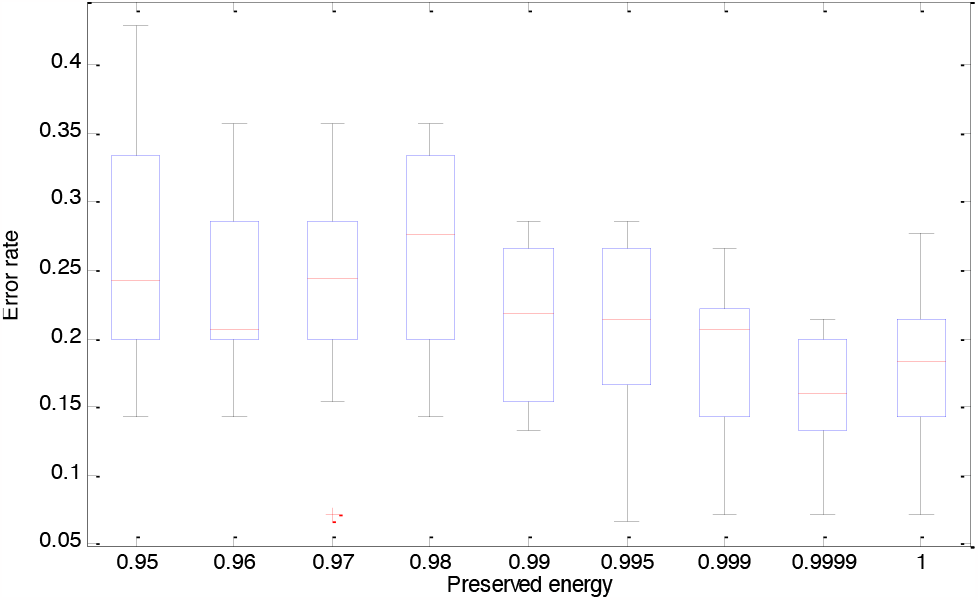
Classification Error Boxplot in 10f-CV.

Median as well as first and third quartiles of the obtained error values are shown in this figure. Horizontal axis is the amount of preserved energy in the PCA. As can be seen in Figure 3, the lowest median and error variance are obtained for preserving 99.99 percent of the energy. Therefore, related dimension to this amount of energy preservation is considered the optimal dimension for applying PCA algorithm [41].

Results of the SRC method using BP, OMP and SL0 sparse coding algorithms are given in Table 2. All the reported results have been obtained using MATLAB 2016 on a core i5 computer with 2.13 GHz processor.

**TABLE II.**
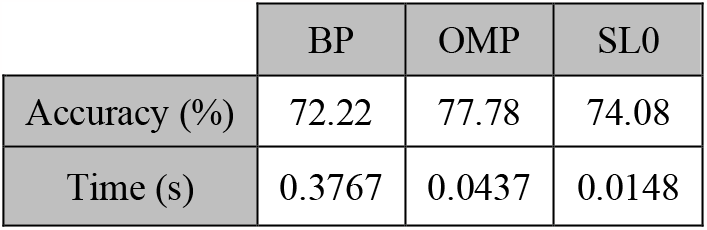
SRC Results.

As the results show the best accuracy is obtained using the OMP while the SL0 algorithm is computationally more efficient. The BP algorithm has the lowest performance.

As mentioned in section 4.2., due to the use of *ℓ*^2^ - norm minimization, the CRC method is faster than the other sparse representation classification methods. Applying this method leads to 66.67% classification accuracy in 4.45 × 10^−4^ seconds. The KNN-SRC method also leads to 55.56% accuracy in 0.0099 seconds for *K* = 26. The optimal value of K was also determined using the10f-CV method.

Classification results of MSRC using different sparse coding algorithms are also summarized in Table 3. In this application, a metasample is equivalent to a linear combination of gene expression profiles from several samples which collects their intrinsic attributes and summarizes the gene expression pattern [21]. In order to evaluate the performance of this method, after dimension reduction by PCA, metasamples are determined using SVD for each class separately, and the final dictionary is constructed by concatenating the obtained metasamples.

**TABLE III.** MSRC Results.

|  | BP | OMP | SL0 |
| --- | --- | --- | --- |
| Accuracy (%) | 79.63 | 79.63 | 79.63 |
| Time (s) | 0.1173 | 0.0445 | 0.0363 |

Due to the unbalanced number of training samples in different classes, the classifier will be biased to the classes that have more training samples if the number of extracted metasamples for each class is considered equal to the number of its training samples [21]. In order to avoid this problem, the number of metasamples in different classes is considered identical. Based on our experiments, the optimal number of metasamples was equal to 5 which was determined using the 10f-CV approach.

From Table 3, it can be seen that the classification accuracy resulting from all the three algorithms are identical and the main difference lies in the speed of the algorithms in which the minimum time belongs to the SL0 algorithm.

Table 4 also shows the LASRC results.

**TABLE IV.** LASRC Results.

|  | BP | OMP | SL0 |
| --- | --- | --- | --- |
| Accuracy (%) | 77.78 | 79.63 | 79.63 |
| Time (s) | 0.3655 | 0.0342 | 0.0141 |

As Table 4 shows, the SL0 and OMP algorithms lead to higher classification accuracy compared to the BP algorithm. Moreover, these algorithms are much faster. The SL0 is even faster so that the computation time of the OMP is approximately 2.5 times that of the SL0. In order to determine the optimal number of selected atoms from *ℓ*^2^ -norm minimization step, the 10f-CV was used and the best result was obtained by selecting 100 atoms form the original dictionary.

Because of the within class variations in tumor samples, it is expected that the SSC-SRC method leads to good results. Such a method firstly finds the most similar training samples to the test sample and then classifies it using the associated cluster samples. Using BP, OMP and SL0 algorithms, the results of this method are summarized in Table 5.

**TABLE V.** SSC-SRC Results.

|  | BP | OMP | SL0 |
| --- | --- | --- | --- |
| Accuracy (%) | 79.63 | 79.63 | 79.63 |
| Time (s) | 1.4216 | 0.0618 | 0.0232 |

It can be seen that good performances are achieved by applying all three sparse coding algorithms while, as expected, the SL0 is faster than OMP and much faster than the BP one. In the SSC-SRC method, optimal number of clusters is 5 which was determined again by 10f-CV. In this method, only a subset of training samples is passed to the final classification process. Therefore, the possibility that irrelevant samples play a key role in the sparse representation of a test sample is reduced which increases the accuracy of the algorithm compare to the SRC.

Table 6 summarizes the results of different sparse representation-based classifiers. In each method, the best achieved classification accuracy has been reported.

**TABLE VI.** SR-based Classifiers Results.

|  | Accuracy (%) | Time (s) |
| --- | --- | --- |
| SRC | 77.78 | 0.0437 |
| CRC | 66.67 | 0.0004 |
| KNN-SRC | 55.56 | 0.0099 |
| MSRC | 79.63 | 0.0363 |
| LASRC | 79.63 | 0.0141 |
| SSC-SRC | 79.63 | 0.0232 |

The MSRC, LASRC and SSC-SRC outperform the other methods. Although, the overall classification accuracy of these methods is similar, they have different performances for different classes. Figure 4 demonstrates the results of these algorithms for each class separately.

**Fig. 4.**
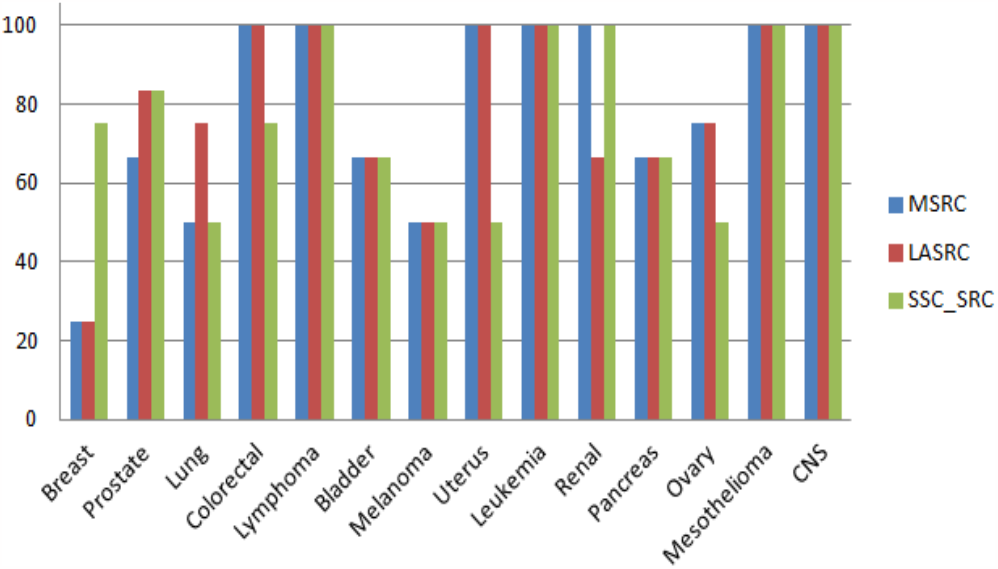
Classification Accuracy for Individual Classes using MSRC, LASRC and SSC-SRC methods.

It can be seen that among the fourteen tumors existing in the database, four tumor classes, namely Lymphoma, Leukemia, Mesothelioma and Central Nervous System (CNS) are perfectly classified (100% accuracy) using all three methods. Three of these four kinds of tumor (Lymphoma, Leukemia and Central Nervous System) have the largest number of training samples compared to the other classes. Melanoma is not classified by any method with an accuracy above 50%. Both Breast and Lung tumors are poorly classified (25% and 50%, respectively) by the two methods. It is observed that the best accuracies are obtained for the classes which have more training samples. This is another confirmation for the impressive role of the number of training samples on the classification performance.

To further investigate the effect of the number of training samples on the classification performance, we perform more experiments using the Leave One Out Cross Validation (LOOCV) method. For this purpose, all the training and test samples are gathered as a pool and one sample is selected as a test sample each time and the other samples are considered the training samples. This process is repeated for all the samples as the test sample. Table 7 shows the results of this experiment using the MSRC, LASRC and SSC-SRC methods.

**TABLE VII.** LOOCV Results.

| Method | Accuracy (%) |
| --- | --- |
| MSRC | 82.83 |
| LASRC | 82.83 |
| SSC-SRC | 83.34 |

These results show that better results are obtained in all cases. Further improvement is achieved for the SSC-SRC method by increasing the number of training samples.

Results of different classification methods applied to 14-Tumors database are shown in Table 8. The SVM and SR results are given in [9]. For each method, the best results have been reported.

**TABLE VIII.** Different Classifiers Results.

| Method | Accuracy (%) |
| --- | --- |
| SVM (OVR) | 75.29 |
| SVM (OVO) | 46.39 |
| SVM (DAG) | 45.1 |
| SVM (WW) | 65.84 |
| SVM (CS) | 75.38 |
| SR | 74.04 |
| SRC | 77.78 |
| SSC-SRC | 79.63 |

From the results in Table 8, it is evident that sparse representation-based classifiers outperform the other methods which often suffer from considerable computational complexity as well.

## 6. Discussion

Our experiments show that the SRC method has a better performance in comparison with the other classifiers. An important advantage of this approach is the lack of need for training which obviates the problem of overfitting to the training data.

In the CRC method, participation of all classes in construction of a dictionary is considered more effective than the sparsity of the coefficients. Therefore, if the number of training samples used in the dictionary is considerably more than the data dimensions, the *ℓ*^2^ -norm will lead to a better performance compared to the *ℓ*^1^ - norm. In the case here, since the number of samples is very small, the weaker performance of the CRC is expected. Moreover, since the *ℓ*^2^ -norm minimization solution is not sparse enough and is spread on almost all the training samples, the least square based methods are highly sensitive to the distribution of the data in different classes. Therefore, imbalanced distribution of data of different classes can be regarded as another influential factor for the poor performance of CRC.

The acceptable performance of MSRC indicates that extracted metasamples from each class deliver a good ability in representation of the class data. The results of the LASRC and SSC-SRC methods confirm that the SRC is improved by these methods. Although these methods select a number of atoms from the original dictionary to determine the sparse representation of a test sample, the criteria used for this purpose are more effective than the one used in the KNN-SRC.

In the LASRC, the *ℓ*^2^ -norm minimization is used and the coefficients with the largest absolute values are selected as the final candidates for concise dictionary formation. In SSC-SRC, samples belonging to the same cluster of a test sample are gathered in order to form the final dictionary. This means that the atom selection processes in these two methods are also based on sparsity concept and are closely related to the sparse representation.

Per class results of the MSRC, LASRC, and SSC-SRC methods indicate that the performance of these methods vary in different classes. Therefore, it can be expected that an appropriate combination of these methods can lead to better results. Also the performance of these methods using the LOOCV evaluation is promising in the sense that better results can be achieved by having more training data. This is especially noteworthy in the SSC-SRC method which shows further improvement in the classification accuracy.

## 7. Conclusions

In this paper, the effectivity of a number of sparse representation-based classification methods on microarray data was studied. Experimental results indicate good performances are achieved using these methods compared to the conventional methods. As shown, even the worst result of these methods that is related to the KNN-SRC presents higher accuracy than some previously adopted methods, such as the SVM (DAG) and SVM (OVO). Among the studied methods, MSRC, LASRC, and SSC-SRC along with the SL0 algorithm lead to the best results. It seems that a better performance can be obtained by a proper fusion of these methods.

Also, sparse representation-based results show that good performances are obtained using a much smaller number of features. In these methods, we have achieved better results by transmitting the data to a lower dimensional space. In our experiments, PCA was used for feature extraction. Moreover, gene selection is a promising approach that addresses the problem of curse of dimensionality and plays an important role in the development of efficient microarray data classification. Due to the fact that only a small number of genes are related to the classification problem, some of the proposed methods try to select the most related genes and eliminate irrelevant and redundant data and noise to improve the classification results [42-46]. Applying feature selection methods before the studied sparse representation-based classifiers can be considered for future works. Evaluation of the performance of the discussed methods on other microarray data is another suggestion for generalizing the obtained results.

## Footnotes

1 Gradient Projection for Sparse Reconstruction

2 Fast Iterative Shrinkage-Thresholding Algorithm

3 Compressive Sampling Matching Pursuit

4 Iterative Hard Thresholding

5 One-Versus-Rest (All)

6 One-Versus-One

7 Directed Acyclic Graph

8 all-at-once method by Weston and Watkins

9 all-at-once method by Crammer and Singer

